# Inoculum effect of antimicrobial peptides

**DOI:** 10.1101/2020.08.21.260620

**Authors:** M. R. Loffredo, F. Savini, S. Bobone, B. Casciaro, H. Franzyk, M. L. Mangoni, L. Stella

## Abstract

The activity of many antibiotics depends on the initial density of cells used in bacteria growth inhibition assays. This phenomenon, termed the inoculum effect, can have important consequences for the therapeutic efficacy of the drugs, since bacterial loads vary by several orders of magnitude in clinically relevant infections. Antimicrobial peptides are a promising class of molecules to fight drug-resistant bacteria, since they act mainly by perturbing the cell membranes rather than by inhibiting intracellular targets. Here we report the first systematic characterization of the inoculum effect for this class of antibacterial compounds. Thirteen peptides (including all-D enantiomers) and peptidomimetics were analyzed by measuring minimum inhibitory concentration values, covering more than 7 orders of magnitude in inoculated cell density. In all cases, we observed a significant inoculum effect for cell densities above 5 × 10^4^ cells/mL, while the active concentrations remained constant (within the micromolar range) for lower densities. In the case of membrane-active peptides, these data can be rationalized by considering a simple model, taking into account peptide-cell association and hypothesizing that a threshold number of cell-bound peptide molecules is required in order to cause a killing effect. The observed effects question the clinical utility of activity and selectivity determinations performed at a fixed, standardized cell density. A routine evaluation of the inoculum dependence of the activity of antimicrobial peptides and peptidomimetics should be considered.

**Significance statement:** Bacterial drug resistance is a crucial threat to global health and antimicrobials with novel mechanisms of action are severely needed. Antimicrobial peptides are natural molecules that kill bacteria mostly by perturbing their membranes and represent promising compounds to fight resistant microbes. Their activity is normally tested under standardized conditions of bacterial density. However, the bacterial load in clinically relevant infections varies by many orders of magnitude. Here we showed that the minimum peptide concentration needed for bacterial killing can vary by more than 100 times with an increase in the density of cells in the initial inoculum of the assay (inoculum effect) These findings question utility of the presently used activity screening assays and our current understanding of antimicrobial peptides.

## Introduction

The minimum inhibitory concentration (MIC) is one of the most common measures for the efficacy of antimicrobial compounds [Andrews 2001, Wiegand 2008]. According to the Clinical and Laboratory Standards Institute (CLSI) and European Committee on Antimicrobial Susceptibility Testing (EUCAST) guidelines, the MIC is the lowest drug concentration that abolishes *in vitro* bacterial growth during a short period (typically 20 hours) when using a standard initial cell density (inoculum) of ∼ 5 × 10^5^ colony-forming units (CFU)/mL in assays performed in broth (with an acceptable range of 2 × 10^5^ to 8 × 10^5^ CFU/mL for CLSI and 3 × 10^5^ to 7 × 10^5^ CFU/mL for EUCAST) [Patel 2015, Weinstein 2019, EUCAST 2003]. The choice of a specific value for the inoculum to be applied is dictated by a need for standardizing the assay in clinical practice [Wiegand 2008], albeit bacterial cell densities in clinically relevant infections *in vivo* range from 1 CFU/mL to 10^9^ CFU/mL (in soft tissue or peritoneal infections) [Kang 2014, König 1998, Bingen 1990].

Soon after the introduction of penicillin for civilian use in the 1940s, it was realized that the active concentration of antibiotics might need to be increased significantly when higher bacterial cell densities are inoculated in the assay medium [Luria 1946, Parker 1946], a phenomenon termed the “inoculum effect” (IE) [Brook 1989]. The IE can be caused by different mechanisms [Karslake 2016, Lenhard 2019], including enzymatic degradation of the drug [Brook 1989, Lenhard 2019], a simple consequence of the number of available drug molecules per cell [Udekwu 2009, zur Wiesch 2015], or altruistic behaviors in which dead cells promote enhanced survival of the remaining bacterial population [Meredith 2015]. Traditionally, an IE has been defined as a ≥8-fold change in MIC when an inoculum 100-fold greater than the CLSI recommendation is used, but recent studies have shown that even subtle differences in inoculum may have a dramatic effect on MIC values [Smith 2018]. The IE is commonly examined by determining the MIC, and in this assay the cell density varies by several orders of magnitude with respect to the initial inoculum, over the many hours in which the bacteria are allowed to grow. However, an IE has been demonstrated also under conditions of constant cell density [Karslake 2016].

High-density bacterial infections, including septic bloodstream and urinary tract infections, endocarditis, and abscesses, are quite prevalent and lack efficacious therapies [Pletzer 2018]. Although some reports have contested the therapeutic relevance of the IE [Craig 2004, Davey 1987], several studies have demonstrated its clinical significance, showing that the MICs determined in the standardized assay were ineffective in the clinical treatment of high density infections [Chapman 1983, Soriano 1988, Soriano 1990, Chuang 1998, Mizunaga 2005, Nannini 2009, Nannini 2013, Karslake 2016, Miller 2018]. In some cases, a concentration 1000 times higher than the MIC is required to cure the infection [Soriano 1988, 1990].

While the IE is well-characterized for traditional antibiotics, little is known about this phenomenon for other antimicrobial compounds. Antimicrobial peptides (AMPs), sometimes referred to as “host-defense peptides”, are produced by all living organisms as a first line of defense against pathogens [Zasloff 2002, Mookherjee 2020, Lazzaro 2020]. These peptides can have many functions [Hancock 2016], but most of them cause direct bactericidal effects that typically involve perturbation of the membrane integrity of microbial cells [Lazzaro 2020, Loffredo 2017]. The majority of known AMPs are short and amphipathic and cationic peptides, capable of binding selectively to the anionic membranes of bacterial cells [Bobone 2019, Vaezi 2020, Paterson 2017, Bobone 2013]. Most AMPs accumulate on the outer leaflet of cell membranes, thereby perturbing their surface tension [Paterson 2017, Orioni 2009]. When a threshold of membrane-bound molecules is reached, the stress is released by the formation of pores or other membrane defects. This mechanism of action has been termed the “carpet” model [Gazit 1996, Bocchinfuso 2009], and it makes development of bacterial resistance particularly difficult [Wimley 2010, Fox 2013, Mookherjee 2020]. Therefore, AMPs represent promising lead compounds in the fight against multidrug-resistant bacteria [Mookherjee 2020, Koo 2019, Fox 2013], which constitute a dramatically increasing worldwide threat [Ventola 2015], and several peptides are undergoing clinical trials [Koo 2019].

Considering their characteristic mechanism of action as compared to that of commonly used antibiotics, the existence of a pronounced IE is not obvious in the case of AMPs. Surprisingly, the IE within this class of molecules has been investigated only in a handful of studies [Savini 2018], which are summarized in Table 1.

**Table 1.** Literature studies of IE for AMPs.

| Peptide | Bacteria | Experiment | Inocula<br>(CFU/mL) | IE* | Reference |
| --- | --- | --- | --- | --- | --- |
| Lactoferricin B | *E. coli* | MBC | $3.5 \times 10^4$ - $3.5 \times 10^8$ | 20 | Jones 1994 |
| Gramicidin S | *S. aureus* | MIC | $10^5$, $10^8$ (for *S. aureus*) | 2-8 | Hartman 2010 |
| PGLA | *E. coli* | | $10^5$, $10^{10}$ (for *E. coli*) | | |
| Pexiganan | *E. coli* | MIC | $5 \times 10^0$ - $5 \times 10^8$ | 100 | Jepson 2016 |
| Dansylated<br>PMAP23 | *E. coli* | MBC | $5 \times 10^4$ - $5 \times 10^8$ | 7 | Savini 2017 |
| LL-37 | *E. coli* | MIC | $5 \times 10^5$ - $2 \times 10^7$ | 10 | Snoussi 2018 |

\* IE indicates the fold increase in MIC or MBC in the inoculum range investigated.

In some of these reports [Jones 1994, Savini 2017] the minimum bactericidal concentrations (MBCs; i.e. the minimum drug concentration killing more than 99.9% of the original bacterial cells [Lorian 2005]), rather than the MICs, were measured. While Jones [1994] determined the MBC under normal growth conditions, in our previous report [Savini 2017] we used a minimal medium that ensured a constant cell density. Different media were used also for the MIC assays: salt-free Luria broth [Hartman 2010], Müller Hinton broth (MHB) [Jepson 2016], or MOPS-based rich defined media (RDM) [Snoussi 2018]. In another study, concerning the activity of MSI 94 against *P. aeruginosa* [Levison 1993], quantitative MIC or MBC determinations were not performed, and hence it is not included in Table 1. However, the time-kill curves obtained at different cell densities were indicative of a significant IE.

Hartman [2010] studied only two cell densities (see Table 1), while Snoussi [2018] investigated less than three orders of magnitude for the inoculum. All other studies [Jones 1994, Jepson 2016, Savini 2017] found an interesting trend: while the active concentration generally depends on the inoculum density, it becomes constant when testing below a certain cell density.

Considering the probable clinical relevance of the IE as well as the scarcity and heterogeneity of available data, we performed a systematic investigation on 11 peptides and peptidomimetics (Table 2). For all compounds MIC testing was performed under the same experimental conditions in order to establish whether the IE is a general property of AMPs, and to investigate its possible origin. We measured MIC values for a range covering more than 7 orders of magnitude of inoculum cell densities.

**Table 2.**
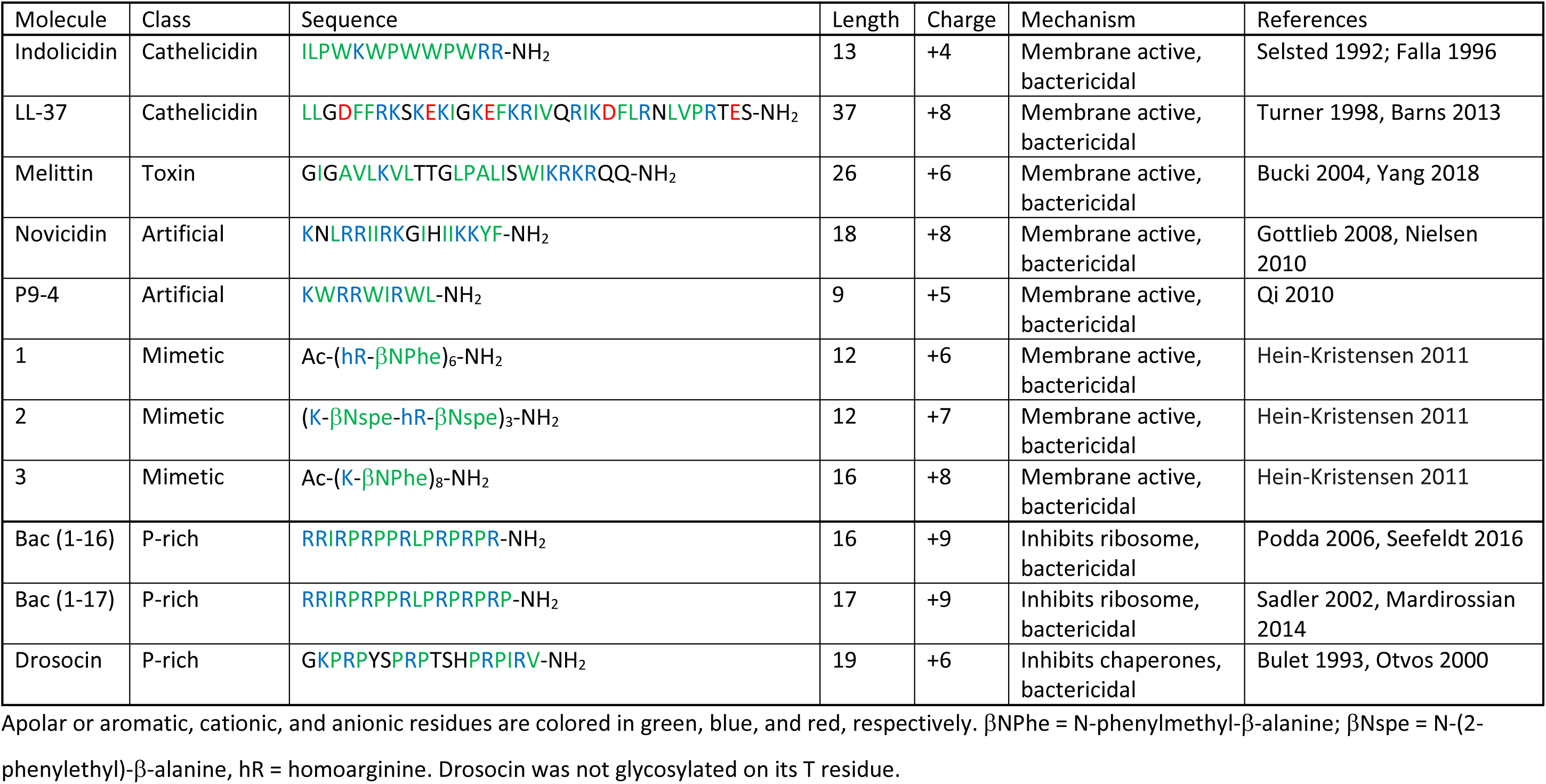
Peptides investigated in the present study and their properties.

As discussed above, for most AMPs, the bacterial membrane is the main target. Among the peptides investigated in the present study, the natural AMPs indolicidin [Selsted 1992; Falla 1996], LL-37 [Turner 1998, Barns 2013], novocidin [Gottlieb 2008, Nielsen 2010], the bee toxin melittin [Bucki 2004, Yang 2018], the artificial peptide P9-4 [Qi 2010] and peptidomimetics 1, 2 and 3 [Hein-Kristensen 2011] all belong to different subclasses of membrane-active antimicrobials that are bactericidal. In principle, upon perturbation of the bacterial membrane some membrane-active AMPs may penetrate into the cell and interact with intracellular targets [Savini 2020, Casciaro 2019]. For instance, indolicidin and LL-37 bind DNA (as many cationic AMPs do), but the role of this phenomenon in the mechanism of bacterial killing is debated [Marchand 2006, Ghosh 2014, Mardirossian 2014, Zhu 2019]. It is also worth mentioning that, in addition to their antimicrobial action, some of these peptides (e.g., LL-37) exert other activities, including immunomodulation and endotoxin neutralisation [Xhindoli 2016]. Other AMPs enter the cell through transporters, without significantly perturbing its membranes, and act on specific intracellular proteins [Cardoso 2019]. As examples of such peptides we included the proline-rich drosocin (in non-glycosylated from) [Bulet 1993, Otvos 2000] as well as fragments 1-16 and 1-17 of bactenecin 7 [Sadler 2002, Podda 2006, Mardirossian 2014, Seefeldt 2016].

## MATERIALS AND METHODS

### Peptides

Melittin was from Sigma-Aldrich, while all other peptides were synthesized on a CEM LibertyTM microwave peptide synthesizer by using microwave-assisted Fmoc-based solid-phase peptide synthesis (SPPS) on a Rink-Amide resin by using previously reported conditions [Hansen 2016]. Peptidomimetics based on the α - peptide/β-peptoid chimeric backbone were prepared by solid-phase synthesis as previously described [Bonke 2008].

### Antimicrobial assays

The MIC values against the Gram-negative bacterium *Escherichia coli* ATCC 25922 were evaluated by using the standard broth microdilution assay [Wiegand 2008] in cation-adjusted Mueller-Hinton II broth (MHB, Becton Dickinson, 212322), as outlined by CLSI [Weinstein 2019, Patel 2015], using 96-well polypropylene microtiter plates (Thermo Scientific Nunc, 267245). *E. coli* was grown in MHB at 37 °C to a mid-log phase. Aliquots of 50 µL of bacterial suspension at different cell densities (dilution from 1 × 10^2^ to 2 × 10^8^) were added to 50 µL MHB containing serial 2-fold dilutions of the peptides (from 64 µM to 0.5 µM) previously prepared in the microplate. After 20 hours incubation at 37 °C, the MIC was determined and defined as the lowest peptide concentration showing no visible growth. MHB diluted 1:2 and 1:10 in water was also used to determine the MIC of the selected peptides, using the same procedure [Merlino 2017, Casciaro 2018]. Experiments on bacterial growth were performed by measuring the optical density at 590 nm in a microplate reader (Infinite M 200; Tecan AG, Switzerland) over 20 h at 37 °C. Untreated bacterial cells were used as the control.

### Peptide adsorption to the multiwell plate surface

Adsorption tests were carried out by using the same 96-well polypropylene microtiter plates (Thermo Scientific Nunc, 267245) used for MIC experiments. Fluorescence data were collected using a Tecan Infinite 200PRO plate reader (Tecan AG, Switzerland). Briefly 300 µL of Buffer A (Phosphate buffer 10 mM, NaCl 140 mM) containing a 2-fold dilution (from 8 to 2 µM) of peptide or tryptophan (used as a control), were added to different wells. The fluorescence signal coming from the center of the well was collected every 10 min for a total time of 1200 min, using λ_ex_ = 280 nm and λ_em_ = 360 nm.

## RESULTS

### 1. Influence of cell density on AMP activity is a universal phenomenon

The IE was evaluated by determining MIC values for inoculum densities ranging from 5 × 10^1^ to 1 × 10^8^ CFU/mL using the microdilution method in cation-adjusted Mueller-Hinton (MH) broth, according to the CLSI protocol (except for the inoculum density) and each measurement was recorded in triplicate.

As illustrated in Figure 1, a significant increase in MIC values (ranging from a factor of 3 to more than 100-fold) was observed for all compounds when applying inoculum densities above the standard value of 5 × 10^5^ CFU/mL. Due to the broad panel of peptides and peptidomimetics investigated, these data confirm the general relevance of the IE for AMPs. Nevertheless, the MIC values did not decrease to 0 when decreasing bacterial cell densities well below the standard value, and in most cases, a clear plateau for the MIC values at declining cell densities was observed. These plateau values (MIC_min_) remained in the micromolar range, and for most peptides no significant variation in MIC was observed in the range between 5 × 10^1^ and 5 × 10^5^ CFU/mL. These findings are in agreement with the few previously available investigations [Jones 1994, Jepson 2016, Savini 2017], and thus infer a general trend for MIC values of AMPs.

**Figure 1:**
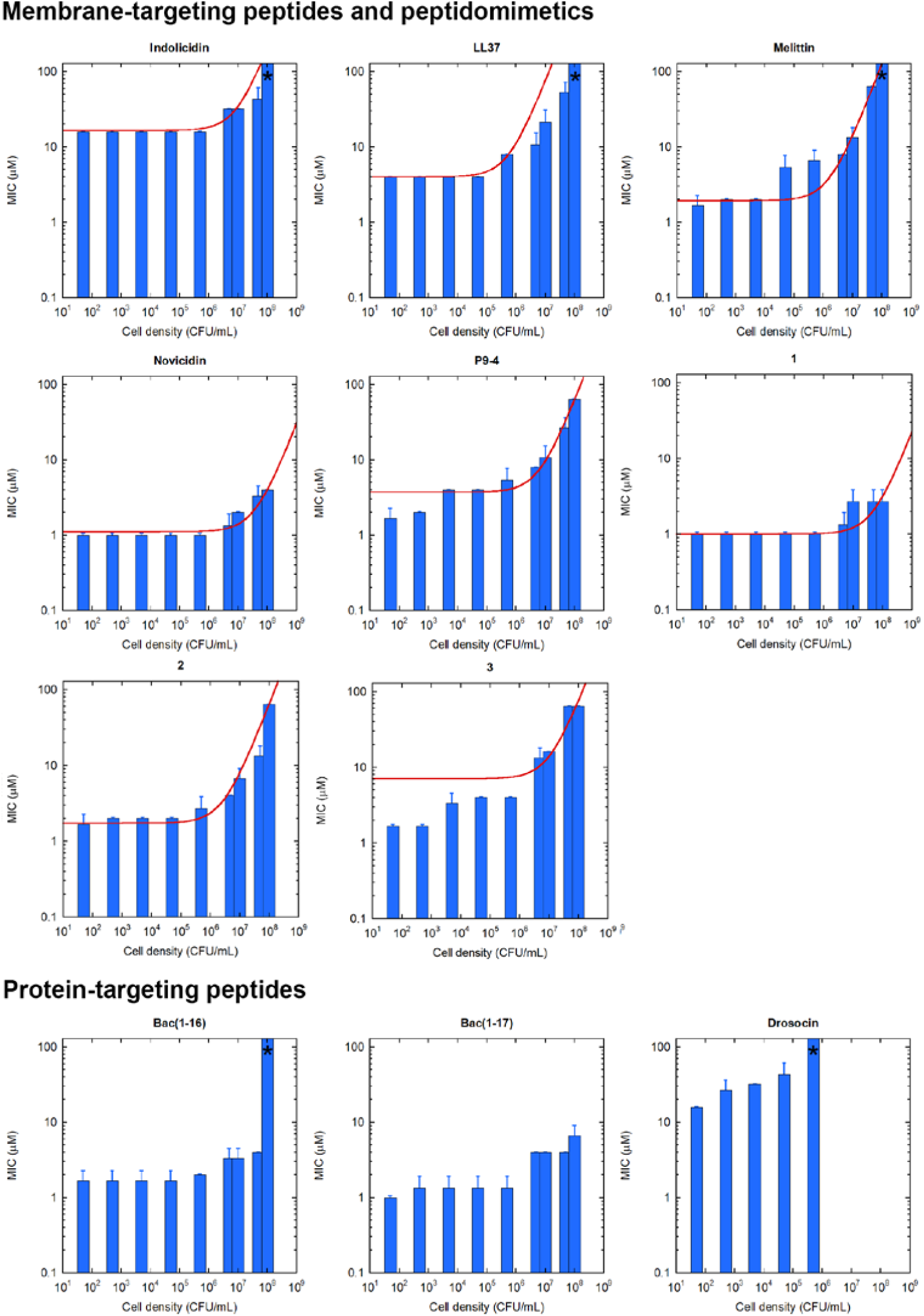
IE for the MIC values of different AMPs and peptidomimetics against *E. coli* ATCC 25922. Each measurement was repeated in triplicate.. Average values and standard deviations (error bars) are reported. An asterisk indicates that the MIC was > 64 µM, since no growth inhibition was observed even at the highest two-fold dilution tested (64 µM). For membrane-active peptides and peptidomimetics, a curve fit with Equation 2 (see discussion) is reported as a red line.

We further determined the time-resolved activity of two selected peptides, indolicidin and Bac(1-16), when tested within the plateau region (i.e., 5 × 10^1^ to 5 × 10^5^ CFU/mL) by following the bacterial growth through measurements of optical density (OD) (λ = 590 nm) for 20 h (Figure 2). Cell cultures at different inocula, not treated with AMPs, grow at the same rate, but reach a measurable OD to different times. At the MIC, no growth was observed. For the membrane-active peptide indolicidin, a concentration equal to half the MIC did not cause any variation in the growth rate, but only a shift in the time at which a significant OD was reached. This finding is consistent with a bactericidal activity and indicates that sub-MIC concentrations kill a fraction of the cells [Jepson 2016]. The remaining ones grow at a similar rate as the initial population, thus requiring a longer time to reach the same final absorbance value, which indicates that they do not differ markedly from the other cells. By contrast, for the AMP with intracellular target, i.e., Bac(1-16), a change in growth rate was observed at 5 × 10^4^ and 5 × 10^5^ CFU/mL.

**Figure 2:**
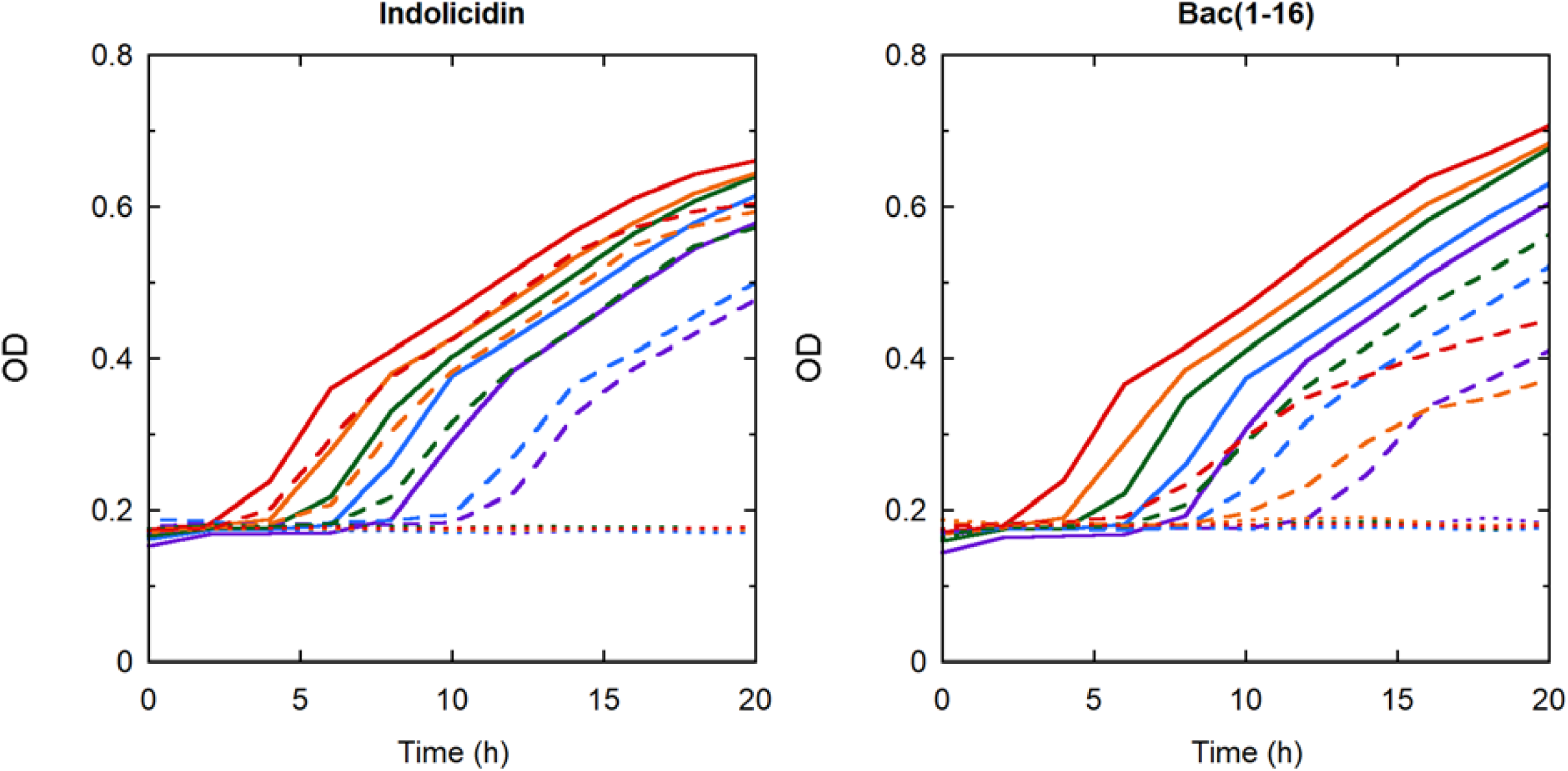
Bacterial growth curves for different inocula (5 × 10^1^, 5 × 10^2^, 5 × 10^3^, 5 × 10^4^ and 5 × 10^5^ CFU/mL, shown in violet, blue, green, orange and red, respectively) of *E. coli* ATCC 25922 in the presence of ½ × MIC (dashed lines) and at MIC (dotted lines) of indolicidin (left) or Bac(1-16) (right) together with the untreated controls (solid lines). OD of the bacterial culture was measured at 590 nm during 20 h incubation at 37 °C. Results are the mean of triplicate samples from a single experiment, representative of three different experiments.

### 2. The inoculum effect is not caused by peptide degradation

For many traditional antibiotics the IE is caused by degradation of the drug by bacterial enzymes [Lenhard 2019]. In the case of AMPs, a common mechanism of bacterial resistance involves production and release of proteases into the extracellular medium, thereby degrading the peptide [Andersson 2016, Bechinger 2017]. To examine whether this phenomenon plays a role also with respect to the IE, enantiomeric peptides (consisting entirely of D-amino acids) not expected to be susceptible to proteases were tested as well. As shown in Figure 3, the MIC of the P9-4 peptide and of its enantiomer exhibited a very similar dependence of cell density. By contrast, in the case of Bac(1-17), which inhibits protein synthesis by binding to the ribosome, the D-enantiomer was much less active, as expected since its mechanism involves a stereospecific interaction with a protein. Noteworthy, peptidomimetics 1, 2 and 3 belong to a compound class previously demonstrated to be resistant to proteolysis, too [Jahnsen 2012]. Based on these data, we can rule out proteolytic degradation as a possible cause of the IE.

**Figure 3:**
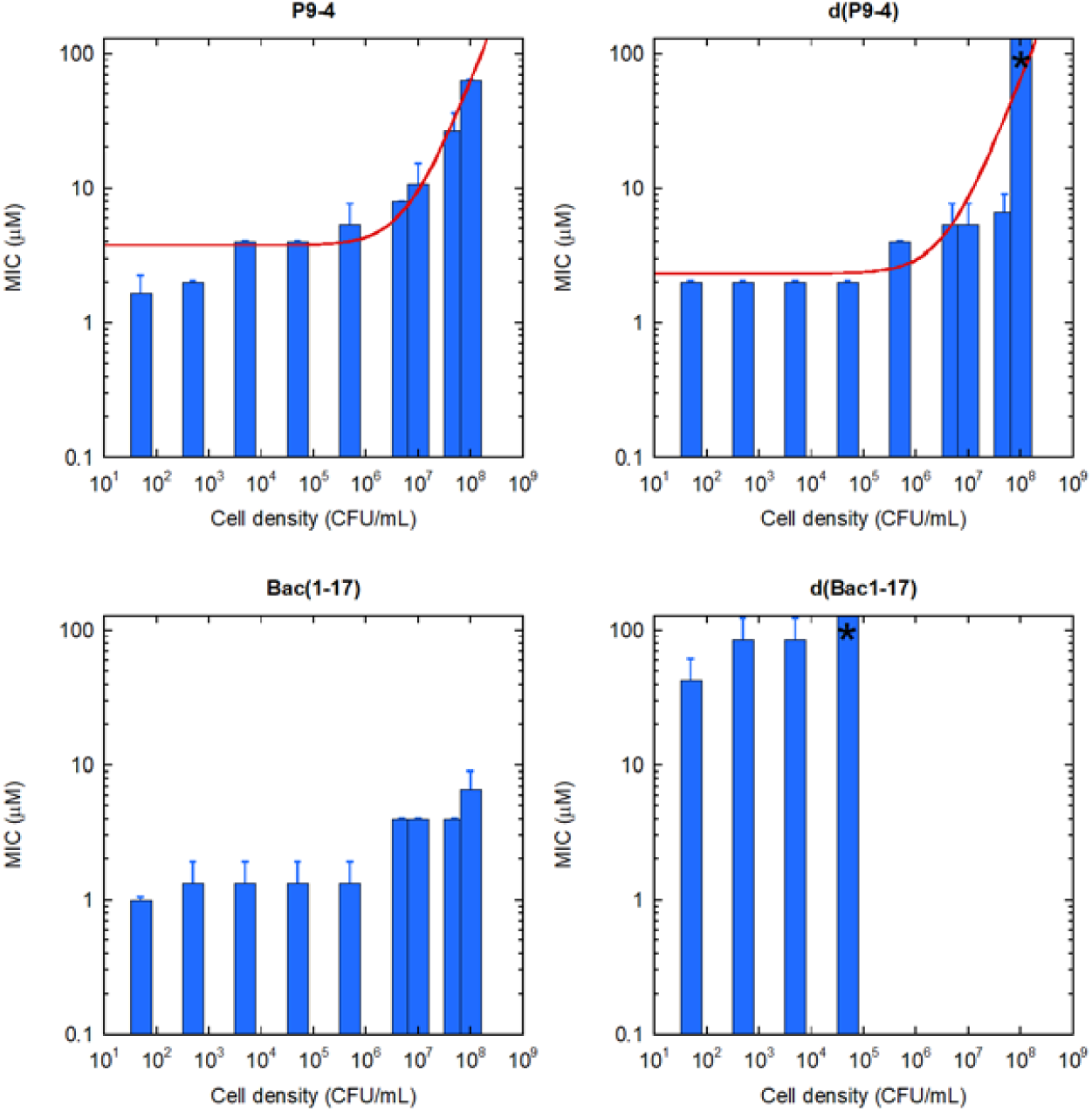
Comparison of the IE for enantiomeric peptides. An asterisk indicates that the MIC was >64 µM, since no growth inhibition was observed even at the highest two-fold dilution tested (64 µM).

### 3. The plateau for MIC values does not arise from experimental artifacts

It has been reported that AMPs can be sequestered by medium components, and their activity is consequently partially lost [Zelezetsky 2006]. It is conceivable that a minimum AMP concentration is needed to saturate the binding sites in the medium components. If this actually was the case, only peptide concentrations exceeding this value would leave some peptide molecules free to interact with the bacteria, resulting in a detectable activity. Therefore, the minimum MIC value observed at vanishing cell densities might in principle correspond to the threshold needed for saturation of the medium components.

To verify the possible role of this propensity for adsorption with respect to the IE, we repeated the MIC determinations in three different dilutions of the MH medium: 100%, 50% and 10%, for the membrane-active peptide LL-37 (Figure 4). This peptide was selected because it has been previously reported that its activity is inhibited by the cell culture medium [Zelezetsky 2006]. LL-37 was found to be more active in the diluted media, possibly as a consequence of reduced peptide sequestration (but also to enhanced bacterial sensitivity due to the lack of nutrients). In any case, its MIC still depended on cell density, and the presence of a plateau at low bacterial loads was conserved. The level of the plateau was only changed by a factor of 4 when the assay was performed in a 10-fold diluted medium. In addition, we have previously observed IE of AMPs and a plateau at low cell densities even in a minimal medium containing only buffer, salts and glucose [Savini 2017]. Therefore, sequestration by medium components can be ruled out as the possible cause of the observed trend.

**Figure 4:**
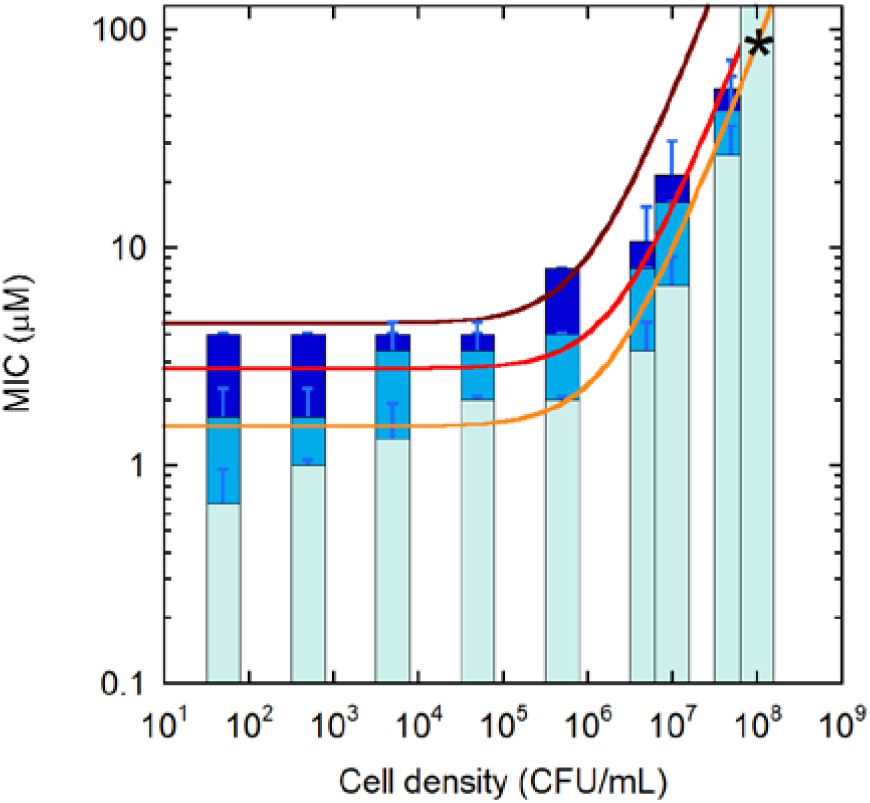
IE of LL-37 against *E. coli* ATCC 25922 after 20 h at 37 °C in 100%, 50% and 10% MH broth (blue, azure and light-blue, respectively).

Considering the amphipathic properties of AMPs, peptide sequestration due to adsorption to the surface of the microwell plate appeared possible [Chico 2003, Goebel-Stengel 2011, Kristensen 2015]. Similarly to peptide sequestration by medium components, a saturation effect in this phenomenon [Kristensen 2015] could explain the plateau value in the MIC at lower cell densities. However, to the best of our knowledge, a real-time measurement of peptide adsorption to container walls over several hours has not been previously reported. To reduce peptide adsorption to the microwell surfaces, we performed our experiments in polypropylene, rather than polystyrene plates [Wiegand 2008, Citterio 2016]. In addition, we tested the effect of peptide adsorption to the microplate walls under the worst possible conditions, i.e., in the absence of bacteria and medium, merely using a simple buffer. Adsorption was followed by utilizing the intrinsic fluorescence of tryptophan residues present in some of the AMPs (i.e., indolicidin, melittin, and d(P9-4)). Recording of the emission signal was focused at the center of the well volume, so that any peptide adsorbed on the walls of the well would not be detected. A control experiment with a tryptophan solution ruled out any photobleaching effects under the experimental conditions used (data not shown). Three concentrations were tested for each peptide: 2, 4 and 8 μM. For indolicidin, all three values are lower than the MIC_min_ (20 μM). By contrast, for melittin and d(P9-4), MIC_min_ (2 μM) coincides with the lowest concentration tested.

For all AMPs, significant adsorption was observed during the 20 h used as incubation time in the MIC determinations (Figure 5). The peptide molecules were adsorbed to the plate walls to different degrees and the effect was concentration-dependent: at 8 µM the adsorbed fractions were about 20%, 30% and 30%, for melittin, d(P9-4) and indolicidin, respectively; these values were much higher at 2 µM (40%, 70%, and 90%), consistently with a saturation effect. However, absorption during the first hour of incubation was negligible at all concentrations tested (Figure 5). Membrane-active AMPs are usually bactericidal, rather than bacteriostatic. As shown in Table 2, a bactericidal mechanism has been demonstrated for all the peptides investigated in the present study. In addition, the bacterial killing by membrane-active AMPs is usually very fast, and all peptides investigated in Figure 5 exert their killing activity in less than an hour [Selsted 1992, Bucki 2004, Qi 2010]. Therefore, peptide adsorption to the multiwell plate appears negligible during the time needed by the peptides to exert their action, and it can be ruled out as the main cause for the plateau in MIC values at vanishing cell densities. Nevertheless, the adsorption to plate material is particularly important to consider when testing very potent AMPs that may display an apparent low activity when applying inappropriate plates for MIC testing [Citterio 2016].

**Figure 5:**
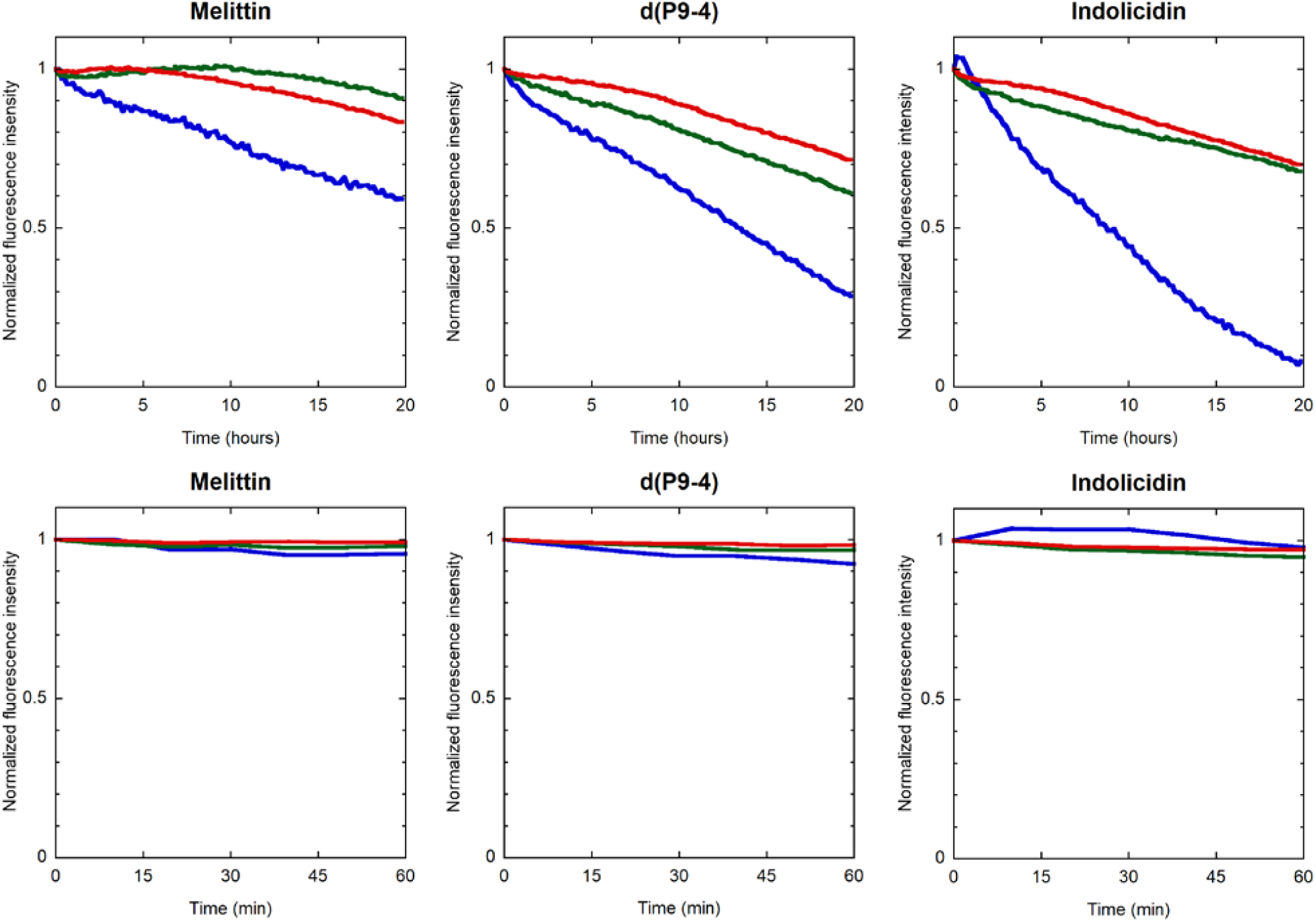
Peptide adsorption to the multiwell plate surface, as determined by the fluorescence in the center of the well volume. Peptide concentrations of 2, 4 and 8 µM are reported in blue, green and red, respectively. Top: 20 hours; bottom: 60 min.

## Discussion

Our data indicate that the IE is a universal property for the activity of AMPs, irrespective of their mechanism of action. In addition, we observed that below a certain peptide-dependent threshold of cell density the MIC becomes constant. This trend is also consistent with the very few previous observations available in the literature, and it seems to be a general characteristic of AMPs.

We ruled out proteolytic degradation, and experimental artefacts as possible causes for the observed behavior. Recently [Savini 2017], we had predicted the observed trend in activity (measured as MIC) vs inoculum density for membrane-active AMPs observed here, based on two simple assumptions that were confirmed experimentally in the case of a PMAP23 analogue [Roversi 2014, Savini 2017, Savini 2018, Savini 2020]. In that case, by performing quantitative studies of AMP interaction with bacterial cells, we have shown that a threshold number of peptides (*T*_*B*_) must be bound to a bacterial cell in order to cause its death [Roversi 2014]. Therefore, the IE may simply arise from the quite obvious fact that a higher inoculum requires a higher peptide concentration to reach this threshold for each cell. Secondly, the plateau reached for the MIC at low cell densities can be explained by the observation that in the presence of bacteria some peptide molecules become associated to the cells, while other remain free in solution. This equilibrium obviously depends on the cell density ([*Bacteria*]). In our studies we have observed that the fraction of cell-bound AMP molecules *f*_*B*_ can be approximately described by the following partition equation [Roversi 2014, Savini 2017]:

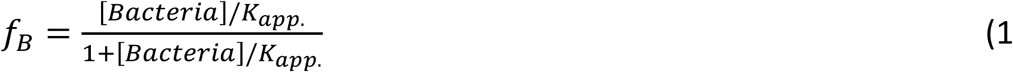

where *K*_*app*._ is an apparent partition constant, corresponding to the cell density at which half of the peptide molecules are cell-bound. As a consequence, the dependence of the active concentration (i.e. the MIC or the MBC, which essentially coincide in the case of a bactericidal mechanism) on cell density can be predicted (see [Savini 2017] for a demonstration):

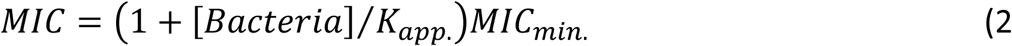

with

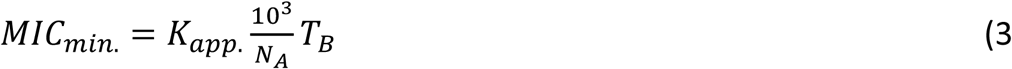

Here *N*_*A*_ is Avogadro’s constant, *MIC and MIC*_*min*._ are expressed in moles/L, *K*_*app*._ while the bacterial cell density ([Bacteria]) is reported in cells/mL, and *T*_*B*_ is expressed in molecules per cell. This linear equation, with a nonzero intercept (*MIC*_*min*_), in a logarithmic cell-density scale corresponds to the trend observed in our experiments, with a plateau reaching *MIC*_*min*._ at inocula significantly lower than *K*_*app*_.

Qualitatively, this behavior can be understood based on the peptide/cell association equilibrium (Figure 6). At cell densities significantly higher than *K*_*app*.._, all the peptide molecules in the sample are associated to bacteria. In this case, the inoculum effect is simply due to the fact that more cells need more peptide molecules to reach *T*_*B*_ that is required for killing. By contrast, in the low cell-density regime ([*Bacteria*] ≪ *K*_*app*._), most of the peptides remain free in solution. In this interval, the partition equilibrium can be approximated by a linear behavior (*f*_*B*_ ≅ [*Bacteria*]/*K*_*app*_), and therefore the fraction of cell-bound peptide molecules decreases proportionally to [*Bacteria*]. As a consequence, these two effects (less cells to kill, and a lower fraction of cell-bound peptide) cancel each other, and the total peptide concentration needed in the sample to kill the bacteria remains essentially constant [Savini 2018].

**Figure 6:**
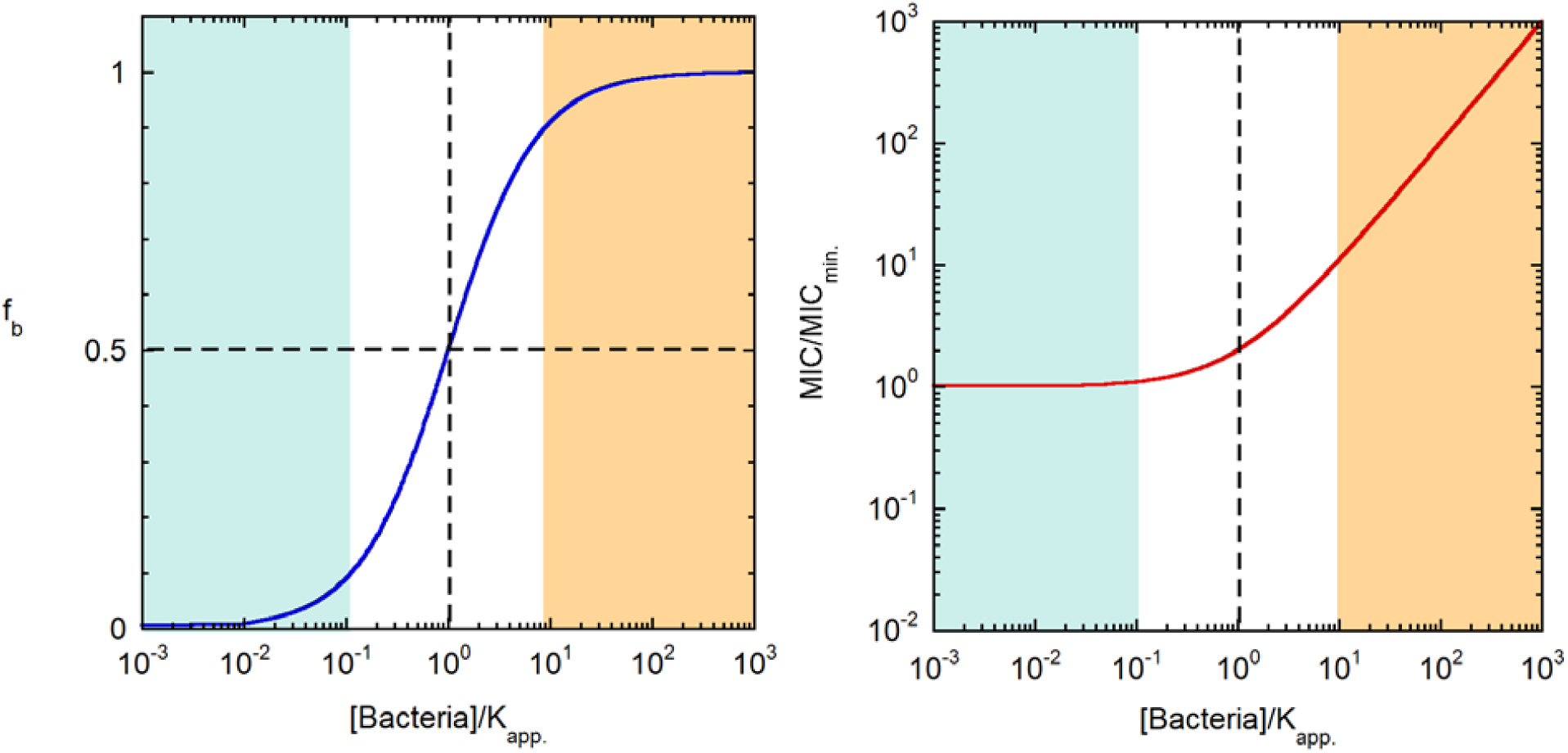
Models for bound peptide/cell and the IE. The fraction of cell-bound peptide molecules (left) and MIC relative to the plateau value at low cell densities (right) are both shown as a function of the inoculum cell density (relative to the apparent partition constant). The blue and orange zones correspond to inocula where the peptides are mostly free in solution, or mostly bound to the cells, respectively.

The MIC data determined in the present study on a large panel of peptides and peptidomimetics were analyzed with the model described above (red curves in Figures 1-3), which describes approximately all the datasets. The parameters obtained with this analysis are reported in Table 3.

**Table 3:**
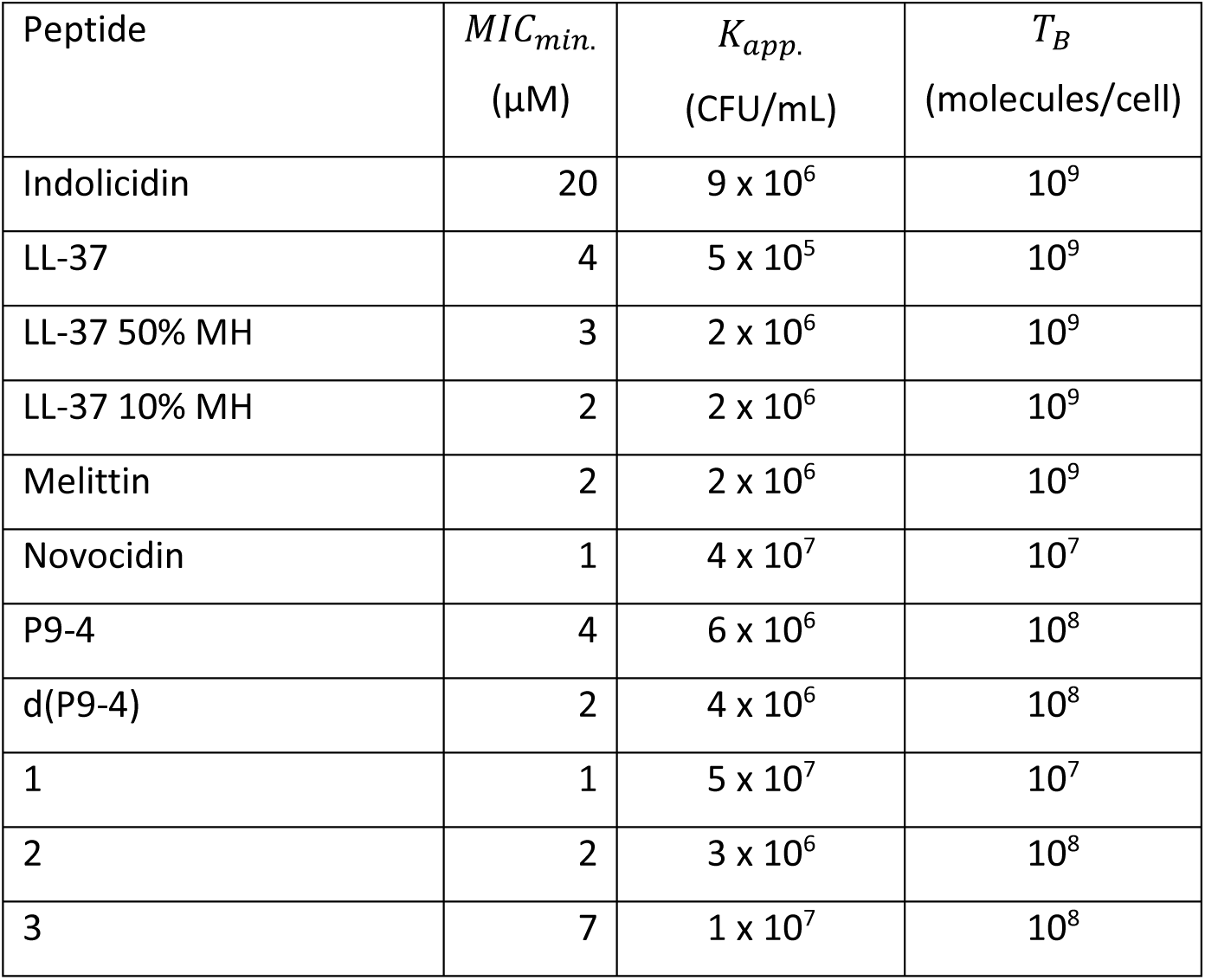
Parameters derived from fitting of the inoculum effect curves (see the main text).

| Peptide | $MIC_{min.}$<br>(μM) | $K_{app.}$<br>(CFU/mL) | $T_B$<br>(molecules/cell) |
| --- | --- | --- | --- |
| Indolicidin | 20 | $9 \times 10^6$ | $10^9$ |
| LL-37 | 4 | $5 \times 10^5$ | $10^9$ |
| LL-37 50% MH | 3 | $2 \times 10^6$ | $10^9$ |
| LL-37 10% MH | 2 | $2 \times 10^6$ | $10^9$ |
| Melittin | 2 | $2 \times 10^6$ | $10^9$ |
| Novocidin | 1 | $4 \times 10^7$ | $10^7$ |
| P9-4 | 4 | $6 \times 10^6$ | $10^8$ |
| d(P9-4) | 2 | $4 \times 10^6$ | $10^8$ |
| 1 | 1 | $5 \times 10^7$ | $10^7$ |
| 2 | 2 | $3 \times 10^6$ | $10^8$ |
| 3 | 7 | $1 \times 10^7$ | $10^8$ |

*MIC*_*min*_ provides a measure of the plateau value of the MIC at vanishing cell densities. All values were in the 1-10 µM range, showing that micromolar AMP concentrations are necessary for antimicrobial activity, even when the bacterial load is extremely low.

The apparent dissociation constant provides a measure of the cell density above which the MIC starts to increase with bacterial counts (more specifically, based on Equation 1, *K*_*app*_ is the cell density for which the MIC is twice *MIC*_*min*._) (Equation 2). The *K*_*app*_ values were within the range 5 ×10^5^ − 5 × 10^7^ CFU/mL. It is worth mentioning that for the PMAP23 labeled analogue the *K*_*app*_ derived from similar IE data was consistent with the *K*_*app*_ measured directly in a binding experiment to *E. coli* cells (1.8 × 10^8^ cells/mL, in that case) [Savini 2017]. Starr [2016] reported apparent binding constants for the artificial AMP ARVA and *E. coli* in the range 5 × 10^6^ − 5 × 10^7^ CFU/mL (depending on peptide concentration).

From the *MIC*_*min*_ and *K*_*app*_ values, according to the model, an estimate for the threshold of bound peptide molecules per cell needed for bacterial killing can be obtained (i.e., *T*_*B*_; Table 3), which determines the strength of the IE, since it defines the slope of the curve (Equations 2 and 3). These values ranged from millions to billions of molecules per cell, confirming our previous conclusion that a huge accumulation of peptide on each cell is required to achieve an antibacterial effect [Roversi 2014, Savini 2017, Savini 2018, Savini 2020]. These numbers are in agreement also with the few previously reported data for the threshold of bound AMP molecules needed for bacterial killing [Steiner 1988, Tran 2002, Roversi 2014, Starr 2016, Savini 2020], which all are within the range of 10^6^-10^8^ molecules/cell (for a review of these data, see [Savini 2018]). However, simple calculations indicate that these numbers correspond to a complete coverage of all the bacterial membranes, or even exceed this limit [Roversi 2014, Savini 2017, Savini 2020]. Therefore, binding to cellular components other than the bacterial membranes is likely. In studies by Jepson [2016], Snoussi [2018] and Wu [2019] it was shown that lysed cells are capable of sequestering a large number of peptide molecules. We have recently measured the association of an AMP to lysed cells, which proved to bind the peptide 10 times stronger than live bacteria [Savini 2020]. Finally, microscopy experiments [Zhu 2019] provided a mechanistic explanation for these observations: once the membranes are permeabilized, bacteria can enter the cell and molecules (e.g., DNA) in the intracellular compartment become accessible for binding. These effects could contribute to the IE by reducing the free peptide concentration available for bacterial killing [Jepson 2016, Snoussi 2018 and Wu 2019], and thus lead to an overestimation of the *T*_*B*_ value. Altruistic behaviors and collective tolerance are common responses in bacteria exposed to antibiotic action [Meredith 2015]. However, these phenomena can come into play only after the bacterial membranes are permeabilized, which still requires a threshold of bound peptides per cell to be reached [Wu 2019].

It is interesting to note that considerations, extremely similar to those presented here, regarding a drug-target binding equilibrium and a threshold number of bound drug molecules per cell to cause killing or growth inhibition, have been demonstrated to be valid also for the IE of traditional antibiotics (e.g., oxacillin, ciprofloxacin or gentamicin) including ionophores such as vancomycin [Udekwu 2009, zur Wiesch 2015]. In addition, a plateau for the MIC values for inocula lower than 10^5^ - 10^6^ has been observed also for traditional antibiotics, irrespective of their mode of action [Udekwu 2009, Artemova 2015, Postek 2018], and this limiting value has by some authors been termed the single-cell MIC [Artemova 2015, Postek 2018]. Therefore, the relevance of the conclusions reached in this study may extend beyond the class of membrane-active AMPs. Indeed, we have observed the same trend for peptidomimetics perturbing the bacterial membrane, and for AMPs inhibiting intracellular targets.

Alternative explanations for the observed trends are of course possible and cannot be ruled out. For instance, high bacterial densities can lead to altered physiological state, gene expression, metabolic activity, age structure, persistence, to the accumulation of metabolic by-products and changes in local pH [Udekwu 2009, Pletzer 2018]. In principle, these effects could reduce the efficacy of AMPs. However, cell density per se appears to be sufficient for explaining the IE. Furthermore, the specific mechanism might be marginally relevant as compared to the consequences of the effect for the appropriate selection of leads to undergo preclinical testing, and for the future clinical application.

Our findings question the suitability of the standard protocol for determining the MIC at a single cell density. The use of a range of cell densities provides a much broader characterization of the efficacy of a peptide (or antibiotic). For instance, our data showed that the activity of different peptides are affected by the cell density to a variable degree. Therefore, a comparison of the activities of various AMPs and analogues should take the IE into account.

The standard inoculum used in MIC assays (i.e., 5 × 10^5^ CFU/mL) falls in the range of *K*_*app*._ values. Since this parameter determines the cell density for which the IE becomes relevant, small errors in the inoculum might affect the MIC values significantly (and to a different extent for different peptides) [Smith 2018]. For traditional antibiotics, it has been suggested that determining the MIC at a lower cell density, where the IE is negligible for all the molecules being characterized, might provide a more robust measurement [Artemova 2015]. Based on the data in Table 3, this condition would correspond to 5 × 10^4^ CFU/mL for AMPs.

We have shown that cell-density affects also the lytic activity of AMPs against erythrocytes [Savini 2017]. This finding supports a general mechanism as the basis for the effect. In addition, it indicates that a complete characterization of the selectivity of AMPs should consider activity and toxicity at different cell concentrations. The selectivity does not have a single value but depends on the density of microbial and host cells [Savini 2017, Savini 2018, Bobone 2019].

Finally, our data indicate that, even at vanishing cell densities, the active concentrations of AMPs do not decrease below the µM range. Such concentrations are reached physiologically in the granules of leukocytes [Hancock 2016] on the skin of frogs [Mangoni 2016] and in the hemolymph of infected insects [Lemaitre 2007]. In addition, several AMPs are normally present in the organism, and they act together, often exhibiting synergism [Mangoni 2009]. Incidentally, the plateau value in the active concentration at low cell densities provides also an explanation for the observation that in some infected insects the production of AMPs is not dependent on the bacterial load, and is maintained for a long time after the infection [Makarova 2016]. However, where the physiological concentrations of AMPs are lower than micromolar, other functions, particularly immunomodulation, might be more important than direct bacterial killing [Hancock 2016].

## Acknowledgements

This work was supported by the Italian Ministry of Education, University and Research (MIUR, grant 667 PRIN 20157WW5EH_007, to L.S.) and Sapienza University of Rome (grant RM11816436113D8A to MLM).

## References

Andersson, D. I., Hughes, D., & Kubicek-Sutherland, J. Z. (2016). Mechanisms and consequences of bacterial resistance to antimicrobial peptides. Drug Resistance Updates, 26, 43–57.

Andrews, J. M. (2001). Determination of minimum inhibitory concentrations. Journal of antimicrobial Chemotherapy, 48(suppl_1), 5–16.

Artemova, T., Gerardin, Y., Dudley, C., Vega, N. M., & Gore, J. (2015). Isolated cell behavior drives the evolution of antibiotic resistance. Molecular systems biology, 11(7).

Barns, K. J., & Weisshaar, J. C. (2013). Real-time attack of LL-37 on single Bacillus subtilis cells. Biochimica et Biophysica Acta (BBA)-Biomembranes, 1828(6), 1511–1520.

Bechinger, B., & Gorr, S. U. (2017). Antimicrobial peptides: mechanisms of action and resistance. Journal of dental research, 96(3), 254–260.

Bingen, E., Lambert-Zechovsky, N., Mariani-Kurkdjian, P., Doit, C., Aujard, Y., Fournerie, F., & Mathieu, H. (1990). Bacterial counts in cerebrospinal fluid of children with meningitis. European Journal of Clinical Microbiology and Infectious Diseases, 9(4), 278–281.

Bobone, S., & Stella, L. (2019). Selectivity of Antimicrobial Peptides: A Complex Interplay of Multiple Equilibria. In Antimicrobial Peptides (pp. 175–214). Springer, Singapore.

Bocchinfuso, G., Palleschi, A., Orioni, B., Grande, G., Formaggio, F., Toniolo, C., … & Stella, L. (2009). Different mechanisms of action of antimicrobial peptides: insights from fluorescence spectroscopy experiments and molecular dynamics simulations. Journal of Peptide Science, 15(9), 550–558.

Bobone, S., Bocchinfuso, G., Park, Y., Palleschi, A., Hahm, K. S., & Stella, L. (2013). The importance of being kinked: role of Pro residues in the selectivity of the helical antimicrobial peptide P5. Journal of Peptide Science, 19(12), 758–769.

Bonke, G., Vedel, L., Witt, M., Jaroszewski, J. W., Olsen, C. A., & Franzyk, H. (2008). Dimeric building blocks for solid-phase synthesis of α-peptide-β-peptoid chimeras. Synthesis, 2008(15), 2381–2390.

Brook, I. (1989). Inoculum effect. Reviews of Infectious Diseases, 11(3), 361–368.

Bucki, R., Pastore, J. J., Randhawa, P., Vegners, R., Weiner, D. J., & Janmey, P. A. (2004). Antibacterial activities of rhodamine B-conjugated gelsolin-derived peptides compared to those of the antimicrobial peptides cathelicidin LL-37, magainin II, and melittin. Antimicrobial Agents and Chemotherapy, 48(5), 1526–1533.

Bulet, P., Dimarcq, J. L., Hetru, C., Lagueux, M., Charlet, M., Hegy, G., … & Hoffmann, J. A. (1993). A novel inducible antibacterial peptide of Drosophila carries an O-glycosylated substitution. Journal of Biological Chemistry, 268(20), 14893–14897.

Cardoso, M. H., Meneguetti, B. T., Costa, B. O., Buccini, D. F., Oshiro, K. G., Preza, S. L., … & Franco, O. L. (2019). Non-Lytic Antibacterial Peptides That Translocate Through Bacterial Membranes to Act on Intracellular Targets. International Journal of Molecular Sciences, 20(19), 4877.

Casciaro, B., Lin, Q., Afonin, S., Loffredo, M. R., de Turris, V., Middel, V., … & Mangoni, M. L. (2019). Inhibition of Pseudomonas aeruginosa biofilm formation and expression of virulence genes by selective epimerization in the peptide Esculentin-1a (1-21) NH 2. The FEBS journal, 286(19), 3874–3891.

Casciaro, B., Loffredo, M. R., Luca, V., Verrusio, W., Cacciafesta, M., & Mangoni, M. L. (2018). Esculentin-1a derived antipseudomonal peptides: limited induction of resistance and synergy with aztreonam. Protein and Peptide Letters, 25(12), 1155–1162.

Chapman, S. W., & Steigbigel, R. T. (1983). Staphylococcal β-lactamase and efficacy of β-lactam antibiotics: in vitro and in vivo evaluation. Journal of Infectious Diseases, 147(6), 1078–1089.

Chuang, Y. C., Ko, W. C., Wang, S. T., Liu, J. W., Kuo, C. F., Wu, J. J., & Huang, K. Y. (1998). Minocycline and cefotaxime in the treatment of experimental murine Vibrio vulnificus infection. Antimicrobial Agents and Chemotherapy, 42(6), 1319–1322.

Chico, D. E., Given, R. L., & Miller, B. T. (2003). Binding of cationic cell-permeable peptides to plastic and glass. Peptides, 24(1), 3–9.

Citterio, L., Franzyk, H., Palarasah, Y., Andersen, T. E., Mateiu, R. V., & Gram, L. (2016). Improved in vitro evaluation of novel antimicrobials: potential synergy between human plasma and antibacterial peptidomimetics, AMPs and antibiotics against human pathogenic bacteria. Research in Microbiology, 167(2), 72–82.

Craig, W. A., Bhavnani, S. M., & Ambrose, P. G. (2004). The inoculum effect: fact or artifact?

Davey, P. G., & Barza, M. (1987). The inoculum effect with gram-negative bacteria in vitro and in vivo. Journal of Antimicrobial Chemotherapy, 20(5), 639–644.

European Committee for Antimicrobial Susceptibility Testing (EUCAST) of the European Society of Clinical Microbiology and Infectious Diseases (ESCMID). (2003). Determination of minimum inhibitory concentrations (MICs) of antibacterial agents by broth dilution. Clinical Microbiology and Infection, 9(8), ix–xv.

Falla, T. J., Karunaratne, D. N., & Hancock, R. E. (1996). Mode of action of the antimicrobial peptide indolicidin. Journal of Biological Chemistry, 271(32), 19298–19303.

Fox, J. L. (2013). Antimicrobial peptides stage a comeback (vol 31, pg 379, 2013). Nature Biotechnology, 31(12), 1066–1066.

Gazit, E., Miller, I. R., Biggin, P. C., Sansom, M. S., & Shai, Y. (1996). Structure and orientation of the mammalian antibacterial peptide cecropin P1 within phospholipid membranes. Journal of Molecular Biology, 258(5), 860–870.

Ghosh, A., Kar, R. K., Jana, J., Saha, A., Jana, B., Krishnamoorthy, J., … & Bhunia, A. (2014). Indolicidin targets duplex DNA: structural and mechanistic insight through a combination of spectroscopy and microscopy. ChemMedChem, 9(9), 2052–2058.

Goebel-Stengel, M., Stengel, A., Taché, Y., & Reeve Jr, J. R. (2011). The importance of using the optimal plasticware and glassware in studies involving peptides. Analytical Biochemistry, 414(1), 38–46.

Gottlieb, C. T., Thomsen, L. E., Ingmer, H., Mygind, P. H., Kristensen, H. H., & Gram, L. (2008). Antimicrobial peptides effectively kill a broad spectrum of Listeria monocytogenes and Staphylococcus aureus strains independently of origin, sub-type, or virulence factor expression. BMC Microbiology, 8(1), 205.

Hancock, R. E., Haney, E. F., & Gill, E. E. (2016). The immunology of host defence peptides: beyond antimicrobial activity. Nature Reviews Immunology, 16(5), 321.

Hansen, A. M., Bonke, G., Larsen, C. J., Yavari, N., Nielsen, P. E., & Franzyk, H. (2016). Antibacterial peptide nucleic acid–antimicrobial peptide (PNA–AMP) conjugates: Antisense targeting of fatty acid biosynthesis. Bioconjugate Chemistry, 27(4), 863–867.

Hartmann, M., Berditsch, M., Hawecker, J., Ardakani, M. F., Gerthsen, D., & Ulrich, A. S. (2010). Damage of the bacterial cell envelope by antimicrobial peptides gramicidin S and PGLa as revealed by transmission and scanning electron microscopy. Antimicrobial Agents and Chemotherapy, 54(8), 3132–3142.

Hein-Kristensen, L., Knapp, K. M., Franzyk, H., & Gram, L. (2011). Bacterial membrane activity of α-peptide/β-peptoid chimeras: influence of amino acid composition and chain length on the activity against different bacterial strains. BMC Microbiology, 11(1), 144.

Jahnsen, R. D., Frimodt-Møller, N., & Franzyk, H. (2012). Antimicrobial activity of peptidomimetics against multidrug-resistant Escherichia coli: a comparative study of different backbones. Journal of Medicinal Chemistry, 55(16), 7253–7261.

Jepson, A. K., Schwarz-Linek, J., Ryan, L., Ryadnov, M. G., & Poon, W. C. (2016). What is the ‘minimum inhibitory concentration’(mic) of pexiganan acting on escherichia coli?—a cautionary case study. In Biophysics of Infection (pp. 33–48). Springer, Cham.

Jones, E. M., Smart, A., Bloomberg, G., Burgess, L., & Millar, M. R. (1994). Lactoferricin, a new antimicrobial peptide. Journal of Applied Bacteriology, 77(2), 208–214.

Kang, D. K., Ali, M. M., Zhang, K., Huang, S. S., Peterson, E., Digman, M. A., … & Zhao, W. (2014). Rapid detection of single bacteria in unprocessed blood using Integrated Comprehensive Droplet Digital Detection. Nature Communications, 5(1), 1–10.

Karslake, J., Maltas, J., Brumm, P., & Wood, K. B. (2016). Population density modulates drug inhibition and gives rise to potential bistability of treatment outcomes for bacterial infections. PLoS Computational Biology, 12(10).

König, C., Simmen, H. P., & Blaser, J. (1998). Bacterial concentrations in pus and infected peritoneal fluid--implications for bactericidal activity of antibiotics. Journal of Antimicrobial Chemotherapy, 42(2), 227–232.

Koo, H. B., & Seo, J. (2019). Antimicrobial peptides under clinical investigation. Peptide Science, 111(5), e24122.

Kristensen, K., Henriksen, J. R., & Andresen, T. L. (2015). Adsorption of cationic peptides to solid surfaces of glass and plastic. PLoS One, 10(5), e0122419.

Lazzaro, B. P., Zasloff, M., & Rolff, J. (2020). Antimicrobial peptides: Application informed by evolution. Science, 368(6490).

Lemaitre, B., & Hoffmann, J. (2007). The host defense of Drosophila melanogaster. Annu. Rev. Immunol., 25, 697–743.

Lenhard, J. R., & Bulman, Z. P. (2019). Inoculum effect of β-lactam antibiotics. Journal of Antimicrobial Chemotherapy, 74(10), 2825–2843.

Levison, M. E., Pitsakis, P. G., May, P. L., & Johnson, C. C. (1993). The bactericidal activity of magainins against Pseudomonas aeruginosa and Enterococcus faecium. Journal of Antimicrobial Chemotherapy, 32(4), 577–585.

Loffredo, M. R., Ghosh, A., Harmouche, N., Casciaro, B., Luca, V., Bortolotti, A., … & Mangoni, M. L. (2017). Membrane perturbing activities and structural properties of the frog-skin derived peptide Esculentin-1a (1-21) NH2 and its Diastereomer Esc (1-21)-1c: Correlation with their antipseudomonal and cytotoxic activity. Biochimica et Biophysica Acta (BBA)-Biomembranes, 1859(12), 2327–2339.

Lorian, V. (2005) Antibiotics in Laboratory Medicine, 5th ed. Lippincott Williams & Wilkins: Philadelphia, PA.

Luria, S. E. (1946). A test for penicillin sensitivity and resistance in Staphylococcus. Proceedings of the Society for Experimental Biology and Medicine, 61(1), 46–51.

Makarova, O., Rodriguez-Rojas, A., Eravci, M., Weise, C., Dobson, A., Johnston, P., & Rolff, J. (2016). Antimicrobial defence and persistent infection in insects revisited. Philosophical Transactions of the Royal Society B: Biological Sciences, 371(1695), 20150296.

Mangoni, M. L., & Shai, Y. (2009). Temporins and their synergism against Gram-negative bacteria and in lipopolysaccharide detoxification. Biochimica et Biophysica Acta (BBA)-Biomembranes, 1788(8), 1610–1619.

Mangoni, M. L., Di Grazia, A., Cappiello, F., Casciaro, B., & Luca, V. (2016). Naturally occurring peptides from Rana temporaria: antimicrobial properties and more. Current Topics in Medicinal Chemistry, 16(1), 54–64.

Mardirossian, M., Grzela, R., Giglione, C., Meinnel, T., Gennaro, R., Mergaert, P., & Scocchi, M. (2014). The host antimicrobial peptide Bac71-35 binds to bacterial ribosomal proteins and inhibits protein synthesis. Chemistry & Biology, 21(12), 1639–1647.

Marchand, C., Krajewski, K., Lee, H. F., Antony, S., Johnson, A. A., Amin, R., … & Pommier, Y. (2006). Covalent binding of the natural antimicrobial peptide indolicidin to DNA abasic sites. Nucleic Acids Research, 34(18), 5157–5165.

Meredith, H. R., Srimani, J. K., Lee, A. J., Lopatkin, A. J., & You, L. (2015). Collective antibiotic tolerance: mechanisms, dynamics and intervention. Nature chemical biology, 11(3), 182.

Merlino, F., Carotenuto, A., Casciaro, B., Martora, F., Loffredo, M. R., Di Grazia, A., … & Galdiero, M. (2017). Glycine-replaced derivatives of [Pro3, DLeu9] TL, a temporin L analogue: Evaluation of antimicrobial, cytotoxic and hemolytic activities. European Journal of Medicinal Chemistry, 139, 750–761.

Miller, W. R., Seas, C., Carvajal, L. P., Diaz, L., Echeverri, A. M., Ferro, C., … & Munita, J. M. (2018, June). The cefazolin inoculum effect is associated with increased mortality in methicillin-susceptible Staphylococcus aureus bacteremia. In Open Forum Infectious Diseases (Vol. 5, No. 6, p. ofy123). US: Oxford University Press.

Mizunaga, S., Kamiyama, T., Fukuda, Y., Takahata, M., & Mitsuyama, J. (2005). Influence of inoculum size of Staphylococcus aureus and Pseudomonas aeruginosa on in vitro activities and in vivo efficacy of fluoroquinolones and carbapenems. Journal of Antimicrobial Chemotherapy, 56(1), 91–96.

Mookherjee, N., Anderson, M. A., Haagsman, H. P., & Davidson, D. J. (2020). Antimicrobial host defence peptides: Functions and clinical potential. Nature Reviews Drug Discovery, 1–22.

Nannini, E. C., Stryjewski, M. E., Singh, K. V., Bourgogne, A., Rude, T. H., Corey, G. R., … & Murray, B. E. (2009). Inoculum effect with cefazolin among clinical isolates of methicillin-susceptible Staphylococcus aureus: frequency and possible cause of cefazolin treatment failure. Antimicrobial Agents and Chemotherapy, 53(8), 3437–3441.

Nannini, E. C., Singh, K. V., Arias, C. A., & Murray, B. E. (2013). In vivo effects of cefazolin, daptomycin, and nafcillin in experimental endocarditis with a methicillin-susceptible Staphylococcus aureus strain showing an inoculum effect against cefazolin. Antimicrobial Agents and Chemotherapy, 57(9), 4276–4281.

Nielsen, S. B., & Otzen, D. E. (2010). Impact of the antimicrobial peptide Novicidin on membrane structure and integrity. Journal of Colloid and Interface Science, 345(2), 248–256.

Orioni, B., Bocchinfuso, G., Kim, J. Y., Palleschi, A., Grande, G., Bobone, S., … & Stella, L. (2009). Membrane perturbation by the antimicrobial peptide PMAP-23: a fluorescence and molecular dynamics study. Biochimica et Biophysica Acta (BBA)-Biomembranes, 1788(7), 1523–1533.

Otvos, L., O, I., Rogers, M. E., Consolvo, P. J., Condie, B. A., Lovas, S., … & Blaszczyk-Thurin, M. (2000). Interaction between heat shock proteins and antimicrobial peptides. Biochemistry, 39(46), 14150–14159.

Parker, R. F. (1946). Action of Penicillin on Staphylococcus. Effect of Size of Inoculum on the Test for Sensitivity. Proceedings of the Society for Experimental Biology and Medicine, 63(2), 443–446.

Patel, J. B., Cockerill II, F. R., Bradford, P. A., Eliopulos, G. M., Hindler, J. A., Jenkins, S. G., Lewis II, J. S., Limbago, B., Miller, L. A., Nicolau, D. P., Powell, D. P., Swenson, J. M., Traczewski, M. M., Turnidge, J. D., Weistein, M. P., & Zimmer, B. L. (2015) M07-A10: Methods for Dilution Antimicrobial Susceptibility Tests for Bacteria that Grow Aerobically, Approved Standard, 10th ed. Clinical and Laboratory Standards Institute: Wayne, PA.

Paterson, D. J., Tassieri, M., Reboud, J., Wilson, R., & Cooper, J. M. (2017). Lipid topology and electrostatic interactions underpin lytic activity of linear cationic antimicrobial peptides in membranes. Proceedings of the National Academy of Sciences, 114(40), E8324–E8332.

Pletzer, D., & Hancock, R. E. (2018). Is synergy the key to treating high-density infections?. Future Microbiology, 13(15), 1629–1632.

Podda, E., Benincasa, M., Pacor, S., Micali, F., Mattiuzzo, M., Gennaro, R., & Scocchi, M. (2006). Dual mode of action of Bac7, a proline-rich antibacterial peptide. Biochimica et Biophysica Acta (BBA)-General Subjects, 1760(11), 1732–1740.

Postek, W., Gargulinski, P., Scheler, O., Kaminski, T. S., & Garstecki, P. (2018). Microfluidic screening of antibiotic susceptibility at a single-cell level shows the inoculum effect of cefotaxime on E. coli. Lab on a Chip, 18(23), 3668–3677.

Qi, X., Zhou, C., Li, P., Xu, W., Cao, Y., Ling, H., … & Mu, Y. (2010). Novel short antibacterial and antifungal peptides with low cytotoxicity: efficacy and action mechanisms. Biochemical and Biophysical Research Communications, 398(3), 594–600.

Roversi, D., Luca, V., Aureli, S., Park, Y., Mangoni, M. L., & Stella, L. (2014). How many antimicrobial peptide molecules kill a bacterium? The case of PMAP-23. ACS Chemical Biology, 9(9), 2003–2007.

Sadler, K., Eom, K. D., Yang, J. L., Dimitrova, Y., & Tam, J. P. (2002). Translocating proline-rich peptides from the antimicrobial peptide bactenecin 7. Biochemistry, 41(48), 14150–14157.

Savini, F., Luca, V., Bocedi, A., Massoud, R., Park, Y., Mangoni, M. L., & Stella, L. (2017). Cell-density dependence of host-defense peptide activity and selectivity in the presence of host cells. ACS Chemical Biology, 12(1), 52–56.

Savini, F., Bobone, S., Roversi, D., Mangoni, M. L., & Stella, L. (2018). From liposomes to cells: Filling the gap between physicochemical and microbiological studies of the activity and selectivity of host-defense peptides. Peptide Science, 110(5), e24041.

Savini, F., Loffredo, M. R., Troiano, C., Bobone, S., Malanovic, N., Eichmann, T. O., … & Stella, L. (2020). Binding of an antimicrobial peptide to bacterial cells: Interaction with different species, strains and cellular components. Biochimica et Biophysica Acta (BBA)-Biomembranes, 183291.

Seefeldt, A. C., Graf, M., Pérébaskine, N., Nguyen, F., Arenz, S., Mardirossian, M., … & Innis, C. A. (2016). Structure of the mammalian antimicrobial peptide Bac7 (1–16) bound within the exit tunnel of a bacterial ribosome. Nucleic Acids Research, 44(5), 2429–2438.

Selsted, M. E., Novotny, M. J., Morris, W. L., Tang, Y. Q., Smith, W., & Cullor, J. S. (1992). Indolicidin, a novel bactericidal tridecapeptide amide from neutrophils. Journal of Biological Chemistry, 267(7), 4292–4295.

Smith, K. P., & Kirby, J. E. (2018). The inoculum effect in the era of multidrug resistance: minor differences in inoculum have dramatic effect on MIC determination. Antimicrobial Agents and Chemotherapy, 62(8), e00433–18.

Snoussi, M., Talledo, J. P., Del Rosario, N. A., Mohammadi, S., Ha, B. Y., Košmrlj, A., & Taheri-Araghi, S. (2018). Heterogeneous absorption of antimicrobial peptide LL37 in Escherichia coli cells enhances population survivability. Elife, 7, e38174.

Soriano, F., Santamaria, M., Ponte, C., Castilla, C., & Fernandez-Roblas, R. (1988). In vivo significance of the inoculum effect of antibiotics onEscherichia coli. European Journal of Clinical Microbiology and Infectious Diseases, 7(3), 410–412.

Soriano, F., Ponte, C., Santamaria, M., & Jimenez-Arriero, M. (1990). Relevance of the inoculum effect of antibiotics in the outcome of experimental infections caused by Escherichia coli. Journal of Antimicrobial Chemotherapy, 25(4), 621–627.

Starr, C. G., He, J., & Wimley, W. C. (2016). Host cell interactions are a significant barrier to the clinical utility of peptide antibiotics. ACS Chemical Biology, 11(12), 3391–3399.

Steiner, H., Andreu, D., & Merrifield, R. B. (1988). Binding and action of cecropin and cecropin analogues: antibacterial peptides from insects. Biochimica et Biophysica Acta (BBA)-Biomembranes, 939(2), 260–266.

Tran, D., Tran, P. A., Tang, Y. Q., Yuan, J., Cole, T., & Selsted, M. E. (2002). Homodimeric θ-Defensins from Rhesus macaqueLeukocytes Isolation, Synthesis, Antimicrobial Activities, and Bacterial Binding Properties of the Cyclic Peptides. Journal of Biological Chemistry, 277(5), 3079–3084.

Turner, J., Cho, Y., Dinh, N. N., Waring, A. J., & Lehrer, R. I. (1998). Activities of LL-37, a cathelin-associated antimicrobial peptide of human neutrophils. Antimicrobial Agents and Chemotherapy, 42(9), 2206–2214.

Udekwu, K. I., Parrish, N., Ankomah, P., Baquero, F., & Levin, B. R. (2009). Functional relationship between bacterial cell density and the efficacy of antibiotics. Journal of Antimicrobial Chemotherapy, 63(4), 745–757.

Vaezi, Z., Bortolotti, A., Luca, V., Perilli, G., Mangoni, M. L., Khosravi-Far, R., … & Stella, L. (2020). Aggregation determines the selectivity of membrane-active anticancer and antimicrobial peptides: The case of killerFLIP. Biochimica et Biophysica Acta (BBA)-Biomembranes, 1862(2), 183107.

Ventola, C. L. (2015). The antibiotic resistance crisis: part 1: causes and threats. Pharmacy and Therapeutics, 40(4), 277.

Weinstein, M. P., Patel, J. B., Bobenchik, A. M., Campeau, S., Cullen, S., Galas, M. f., Gold, H., Humphries, R. M., Kirn, T. J., Lewis II, J. S., Limbago, B., Mathers, A. J., Mazzulli, T., Richter, S. S., Satlin, M., Schuetz, A. N., Swenson, J. M., Tamma, P. D. (2019) Performance standards for antimicrobial susceptibility testing, 29th ed., Clinical and Laboratory Standards Institute: Wayne, PA.

Wiegand, I., Hilpert, K., & Hancock, R. E. (2008). Agar and broth dilution methods to determine the minimal inhibitory concentration (MIC) of antimicrobial substances. Nature Protocols, 3(2), 163.

Wimley, W. C. (2010). Describing the mechanism of antimicrobial peptide action with the interfacial activity model. ACS Chemical Biology, 5(10), 905–917.

Wu, F., & Tan, C. (2019). Dead bacterial absorption of antimicrobial peptides underlies collective tolerance. Journal of the Royal Society Interface, 16(151), 20180701.

Xhindoli, D., Pacor, S., Benincasa, M., Scocchi, M., Gennaro, R., & Tossi, A. (2016). The human cathelicidin LL-37—A pore-forming antibacterial peptide and host-cell modulator. Biochimica et Biophysica Acta (BBA)-Biomembranes, 1858(3), 546–566.

Yang, Z., Choi, H., & Weisshaar, J. C. (2018). Melittin-induced permeabilization, re-sealing, and re-permeabilization of E. coli membranes. Biophysical Journal, 114(2), 368–379.

Zasloff, M. (2002). Antimicrobial peptides of multicellular organisms. Nature, 415(6870), 389–395.

Zelezetsky, I., Pontillo, A., Puzzi, L., Antcheva, N., Segat, L., Pacor, S., … & Tossi, A. (2006). Evolution of the Primate Cathelicidin Correlation between Structural Variations and Antimicrobial Activity. Journal of Biological Chemistry, 281(29), 19861–19871.

Zhu, Y., Mohapatra, S., & Weisshaar, J. C. (2019). Rigidification of the Escherichia coli cytoplasm by the human antimicrobial peptide LL-37 revealed by superresolution fluorescence microscopy. Proceedings of the National Academy of Sciences, 116(3), 1017–1026.

Zur Wiesch, P. A., Abel, S., Gkotzis, S., Ocampo, P., Engelstädter, J., Hinkley, T., … & Cohen, T. (2015). Classic reaction kinetics can explain complex patterns of antibiotic action. Science Translational Medicine, 7(287), 287ra73–287ra73.

